# Syntenizer 3000: Synteny-based analysis of orthologous gene groups

**DOI:** 10.1101/618678

**Authors:** Camous Moslemi, Cathrine Kiel Skovbjerg, Sara Moeskjær, Stig Uggerhøj Andersen

## Abstract

**Motivation:** The amorphous nature of genes combined with the prevalence of duplication events makes establishing correct genetic phylogenies challenging.

Since homologous gene groups are traditionally formed on basis of sequence similarity, both orthologs and paralogs are often placed in the same gene group by existing tools. Certain tools such as PoFF take syntenic relationship of genes into consideration when forming gene groups. However, a method to form gene groups consisting of only true syntelogs has not yet been developed.

In order to obtain orthologous gene groups consisting of the most likely syntelogs we need a method to filter out paralogs. If one strain has two or more copies of the same gene in a gene group we want to keep only the true syntelog in the group, and remove the paralogous copies by distinguishing between the two using synteny analysis.

**Results:** We present a novel algorithm for measuring the degree of synteny shared between two genes and successfully disambiguate gene groups. This synteny measure is the basis for a number of other useful functions such as gene neighbourhood visualisation to inspect suspect gene groups, strain visualisation for assessing assembly quality and finding genomic areas of interest, and chromosome/plasmid classification of contigs in partially classified datasets.

**Availability:** The latest version of Syntenizer 3000 can be downloaded from the GitHub repository at https://github.com/kamiboy/Syntenizer3000/

Consult the manual.pdf file in the repository for instructions on how to build and use the tool, as well as a in depth explanation of the algorithms utilised.

## 1 Introduction

The amorphous nature of genetic material poses a major challenge for the process of establishing genetic phylogenies. When Needleman & Wunsch introduced a computationally efficient method to look for not just exact copies, but also somewhat similar amino acid sequences in proteins, their method became an industry standard that is still being used today [Needleman & Wunsch 1970]. Often, however, sequence similarity alone is not sufficient for establishing correct genetic phylogenies. This is the case, for example, when focus shifts from homologous to orthologous or paralogous genes [Fitch 1970]. Orthologous genes had a common origin in an ancestor before a speciation event, while paralogous genes are the result of gene duplications within the same species (**Figure 1**).

**Figure 1.**
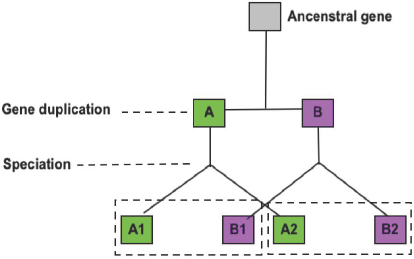
Gene duplication of an ancestral gene results in the paralog pair A and B in the same species. A speciation event then separates these into two different lineages. The gene pairs with the same colour are orthologs in different species, the genes pairs inside the dashed line are paralogs within the same species. Although sequence similarity would lump all four resulting genes into one homology group, only the genes with the same colour are orthologs, a paralog aware grouping would form two separate ortholog pairs, A1 with A2, and B1 with B2.

Studying the evolutionary history of genes by establishing gene homologies, and separating orthologous from paralogous genes [Koonin 2005], has been a topic of great interest and subject to much research over the years. Despite this, no perfect method for automatically forming orthologous groups has been found, and benchmarking different methods is difficult and relies on manually curated datasets [Trachana 2011].

In bacteria, gene duplication is one of the major contributors to ambiguity when establishing gene phylogeny among different strains. Gene duplications are an important driver of bacterial evolution, and the genome of certain bacteria can consist of large portions of related genes [Serres 2009]. Due to sequence similarity it is not easy to establish which of a number of alternate genes in species B are paralogs to or orthologs to a gene in species A.

Ortholog conjecture is one way of distinguishing between such genes based on the notion of orthologs being functionally more similar than paralogs [Altenhoff 2012] since the copied gene tends to drift while the original maintains its original function.

One reason for the differences in drift tendency is that the function of genes is not determined by their sequence alone, but also through the context of their genetic neighbourhood. This can be exemplified by the lactose operon in *Escherichia coli* [Lodish 2000] where a group of genes located near each other all play a role in the same metabolic process. As a consequence, gene function hypotheses can be generated by considering functions of nearby genes [Gerlt & Babbitt 2000].

While gene copies share sequence similarity, the same cannot necessarily be said for their genomic neighbourhoods. The size of duplicated regions is often small in bacteria [Gevers 2004], and therefore the immediate genomic neighbourhood of a duplicate gene will differ from that of the original gene.The flanking regions of each paralog will set them apart when compared to related strains that have the same gene, but have not undergone the same duplication event. If the duplication event is unique to one, or a subset, of strains in a population it should be easy to determine which of the two, or more, copies of a gene is the original (the true syntelog) by cross referencing the other individuals.The only scenario in which this does apply is if a gene duplication event places the paralogs right next to each other in the host genome, in which case their respective flanking regions would be identical (**Figure 2**).

**Figure 2.**
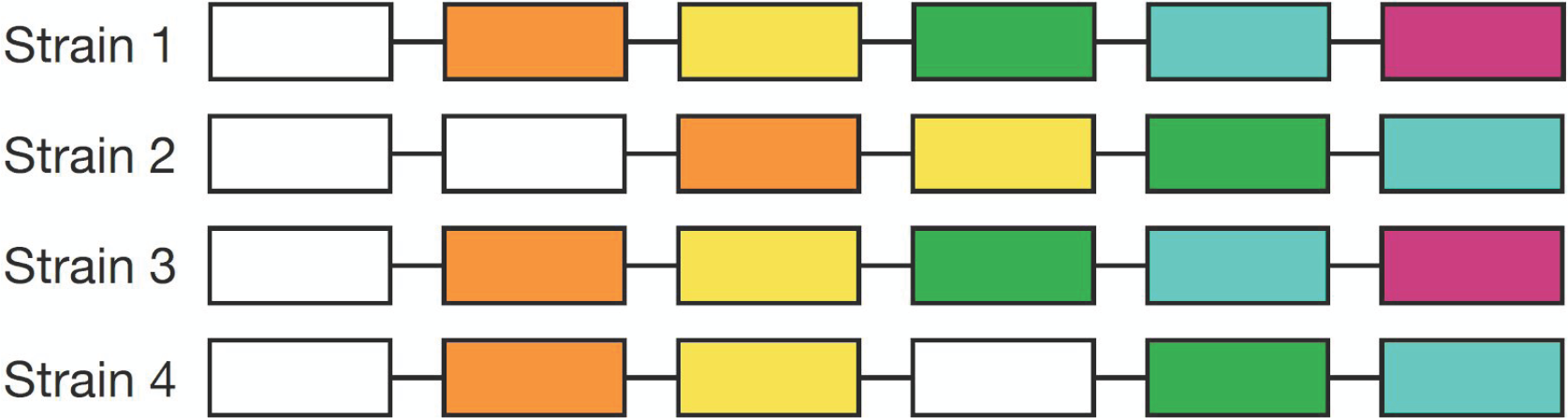
Example of various gene duplication events. The white gene has been duplicated in strain 2 and 4, but not in 1 and 3. In the case of strain 4 it is easy to see which of the two genes is most likely the copy, and which the original by cross referencing neighbouring genes with strain 1 and 3. But in case of strain 2, since the duplicated genes are next to each other, their neighbouring genes are identical, and cross referencing can not reveal which was the copy and which the original.

Thus, in most cases distinguishing between the copy and original is a question of cross referencing the flanking regions of other strains far enough to each side.

Flanking regions can be cross referenced using local alignment, but this is computationally costly. However, if alignment has already been performed on a dataset, for an example via homologous gene group generation, then a much more efficient method involving synteny analysis based on gene groups can be employed, which is what Syntenizer 3000 (S3K) takes advantage of.

## 2 Materials and methods

Since our synteny measuring method relies on pre-existing gene groups these are expected to have already been made using other tools. The sequenced genome should already be assembled, then annotated and have orthologous gene groups generated for it. We have used SPAdes [Bankevich 2012], PROKKA [Seemann 2014], and ProteinOrtho [Lechner 2011] to accomplish these tasks [Cavassim 2019].

### 2.2 Synteny

#### 2.2.1 Defining Synteny

We base our analysis of synteny on the concept of positional orthology as defined in a paper by [Dewey 2011]. The paper defines positional orthology, or synteny, as two genes that are homologous to one another, with neighbouring genes which are also homologous to one another.

If sequence data analysis reveals that three neighbouring genes, A1, B1 and C1, in Strain1 are homologous to the neighbouring genes A2, B2 and C2, respectively, in Strain2, the three homology pairs do not only exhibit orthology but also positional orthology, or synteny.

#### 2.2.2 Measuring Synteny

We base our measurement of synteny on orthologous gene groups. We will regard two genes as being potentially orthologous if they are placed in the same orthology gene group. In effect, our synteny measuring algorithm will only regard genes in terms of the orthologous group that they belong to. Below is the genome of the two strains from Figure 1 regarded in terms of the orthologous group of each gene, rather than the gene itself.

Strain1: ===group1===group2===group3===

Strain2: ===group1===group2===group3===

S3K can assign a score to the strength of synteny between two genes. To help illustrate how the scoring algorithm functions we will use an example of two potentially orthologous genes A1 and A2 that both reside in the same group but come from different strains. We wish to examine synteny between these two genes and for that purpose we will look at their [n] closest neighbouring genes. By neighbouring genes we mean the genes placed directly up- and downstream of the gene in question in the genome of the host organism. In S3K the default value of [n] is set to 40, meaning that 20 upstream genes, and 20 downstream genes will be taken into consideration.

Strain1: ===group7===group6===group5===A1===group2===group3===group4===

Strain2: ===group7===group6===group5===A2===group2===group3===group4===

Above is an example of this concept with an [n]=6 where two genes, A1 and A2, demonstrate perfect synteny for an [n]=6, because each of their 6 neighbouring gene pairs are predicted orthologs. In the course of developing the final synteny scoring algorithm several scoring methods were developed of increasing complexity. Below is an explanation of the two that were kept.

## I. SIMPLE SCORING METHOD

The first method, called SIMPLE, works by finding genes in the [n] neighbourhood of A1 and A2 that belong to the same orthology group. For each match the synteny score between A1 and A2 is increased by 1. So the maximum score possible is 40, indicating perfect synteny in the [n]=40 neighbourhood of both genes, while a score of 0 indicates that no synteny exists between the two genes.

## II. Sophisticated SCORING METHOD

One disadvantage with the SIMPLE scoring method is that the score does not properly reflect gene insertion/deletions and other gene shuffling events.

In order to mitigate this a sophisticated scoring method was developed which first attempts to align the neighbourhoods of two genes akin to the Needleman-Wunsch (NW) algorithm does for DNA sequences [Needleman & Wunsch 1970].

This is the basis of the SOPHISTICATED scoring algorithm which is a modification of the NW algorithm to repurpose it to aligning gene neighbourhoods rather than nucleotides. The NW algorithm is configured with a gap penalty of −0.5, match score of 1 and mismatch score of 0.

Strain1: ===group2===group5===group3===A1===group7===group4===group6===

Strain2: ===group8===group2===group5===A2===group7===group4===group6===

For the above example the algorithm would align the two neighbourhoods as seen below.

Strain1: ===group2===group5===group3===A1===group7===group4===group6===

Strain2: ===group2===group5===---------===A2===group7===group4===group6===

The score assigned would then be 4.5 which is the result of 5 perfect group matches, minus 0.5 penalty for the gap (--------) that needed to be introduced to align the neighbourhoods correctly.

## III. SCORE NORMALISATION

The synteny scoring algorithm described measures synteny for one gene pair. This is only useful for calculating synteny in gene groups with only two members. However, most of orthologous groups in large datasets contain many more genes, with some core groups exceeding the number of strains due to the presence of paralogs. Therefore, a method was developed to assign a synteny score to each gene group by taking into account how each gene in said group shared synteny with each other gene in that group.

Considering an orthologous gene group with three members, the total synteny score of the group is calculated as the sum of synteny scores for each possible unique gene pairing divided by the number of possible unique gene pairings in said group to normalise the scores across groups and make them comparable.

**GroupA**: **[A1], [A2], [A3]**

[**GroupA]**: Score = (*Synteny*(**[A1], [A2]**) + *Synteny*(**[A1], [A3]**) + *Synteny*(**[A2], [A3]**)) **/ 3**

The maximum attainable value of the score normalisation is equal to the size of the neighbourhood [n]=40 for both SIMPLE and SOPHISTICATED scoring methods.

### 2.3 Gene group disambiguation

In order to disambiguate an orthologous gene group which contains a number of co-orthologous genes, the true syntelog must be found among the co-orthologous genes, so all none-true syntelog paralogs can be removed from the gene group, leaving behind a group consisting only of orthologous genes.

Our algorithm for achieving this works in three steps. In the first step all co-orthologous genes are removed and pooled into separate subgroups according to strain, leaving the remaining genes in one large purely orthologous gene group. In the second step all genes in each co-orthologous subgroup are scored against the pure orthologous group using the SOPHISTICATED score. The gene in each co-orthologous subgroup with the highest synteny against the pure orthologous group is chosen as the true syntelog, and is inserted back into the pure ortholog group.

In the third step new gene groups are made from the remaining genes in the co-orthologous subgroups using synteny score to determine which genes are most closely related, while lone left over genes that do not belong in any group are orphaned and discarded.

**Step1:** [A B1 B2 B3 C D E1 E2 F G] =>

**Orthologs**:[A C D F G] + **Co-orthologs**:[B1 B2 B3] + **Co-orthologs**:[E1 E2]

**Step2:** [A C D F G] [B1 B2 B3] [E1 E2] =>

**Orthologs**:[A B2 C D E1 F G] + **Paralogs**:[B1 B3] + **Paralogs**:[E2]

**Step3:** [A B2 C D E1 F G] [B1 B3] [E2] =>

**Orthologs**:[A B2 C D E1 F G] + **Orthologs**:[B1 E2] + **Orphaned**:[B3]

## 4 Results

The datasets used to evaluate the performance of S3K originates from 196 *Rhizobium leguminosarum* strains sampled from clover nodules in various European fields [Cavassim 2019]. Genomic DNA from all strains has been subjected to whole genome shotgun sequencing using 2×250 bp Illumina followed by genome assemblies using SPAdes (v. 3.6.2) [Antipov 2015].

The genomic features of the assemblies for each strain were then annotated using PROKKA [Seemann 2014] which relies on a suite of tools that predict CDS, rRNA, tRNA etc. and predicts CDS function using external curated databases of bacterial proteins.

Orthologous gene groups were obtained using ProteinOrtho [Lechner 2011] using default parameters. A total of 1,424,452 genes in the dataset were located in 6,561 contigs (**Table 1**) with an average number of 33 contigs per strain, meaning that the genomes were fragmented [Izabel 2019].

**Table 1.** Summary statistics generated by S3K for ProteinOrtho gene groups reveals a large number of short contigs (containing less than 41 genes), but only 2% of total genes are located on such contigs.

|  | Before Disambiguation | After Disambiguation |
| --- | --- | --- |
| Number of genes in dataset | 1,424,452 | 1,423,630 |
| Number of contigs in dataset | 6,561 | 6,561 |
| Number of short contigs in dataset | 3,155 (48%) | 3,155 (48%) |
| Number of groups in dataset | 21,198 | 22,128 |
| Avg. number of genes per group | 67.20 | 64.34 |
| Number of co-orthologous genes | 9,428 (0.66%) | 0 |
| Number of groups containing co-orthologs | 1,380 (6.51%) | 0 |
| Number of genes in short contigs | 34,182 (2.40%) | 33,912 (2.38%) |
| Number of groups with short contig genes | 8,351 (39.40%) | 8,723 (39.42%) |

In order to ensure that the high fragmentation would not disrupt the measurement of synteny, the gene group file output from ProteinOrtho as well as the annotated genomes output from PROKKA [Seemann 2014] were input into S3K (**Figure 3**) and used to measure the total average synteny of all gene groups (**Table 2**) by toggling the *“--disambiguategroups”* option.

**Table 2.** Average synteny of subgroups formed of orthologous genes and co-orthologous genes in groups that contain at least one co-orthologous pair.

|  | All groups | Orthologous subgroups | Co-orthologous subgroups |
| --- | --- | --- | --- |
| Total number of genes | 1,424,452 | 316,046 | 16,550 |
| Average synteny before disambiguation | 21.19 | 20.17 | 8.20 |
| Average synteny after disambiguation | 22.52 | - | - |

**Figure 3.**
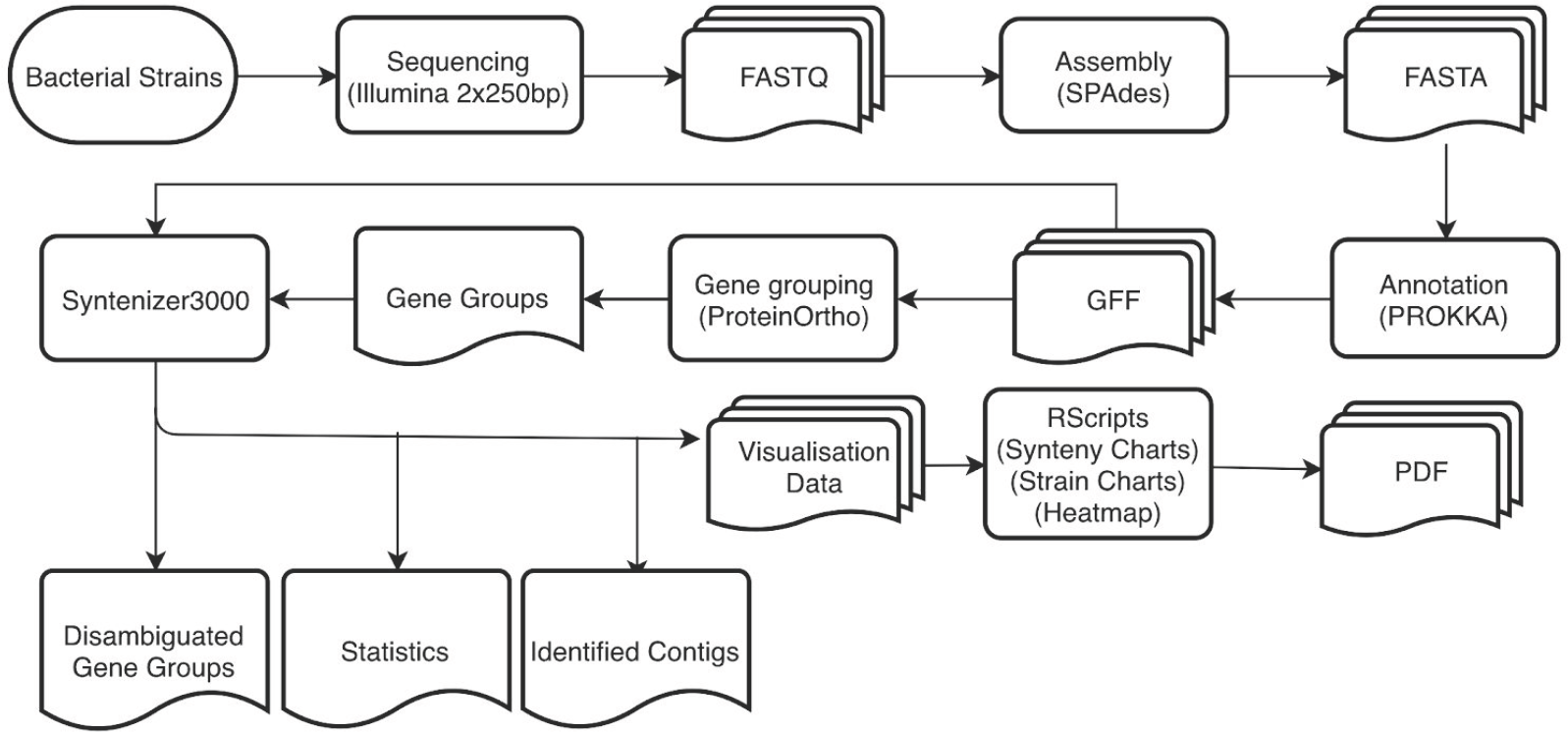
Flowchart of the Syntenizer 3000 toolchain usage.

Fragmentation was revealed not to be an issue given the high average synteny (21.19), plus the low (2.4%) percentage of genes located on contigs with a gene neighbourhood size below 40 (**Table 1**) utilised by the synteny measuring algorithm.

Out of 21,198 groups, approximately 6.51% contained co-orthologs and upon disambiguation an increase of 21.19 to 22.52 in synteny score was observed. The low impact of disambiguation on total average gene group synteny is to be expected as co-orthologous genes only account for 0.66% of the total genome (Table 1).

Two methods of the disambiguation algorithm are available in S3K. Method 1 splits up a gene group, then attempts to reform it based on synteny scores. This approach tends to split some large groups into two or more large groups hinting at the existence of distinct syntenic clusters inside some groups. Method 2 is co-ortholog aware, and designed only to retain the most likely true syntelog version of co-orthologous genes within the group, while attempting to form new groups out of the removed co-orthologs. A gene group synteny visualiser was built into S3K (**Figure 4**).

**Figure 4.**
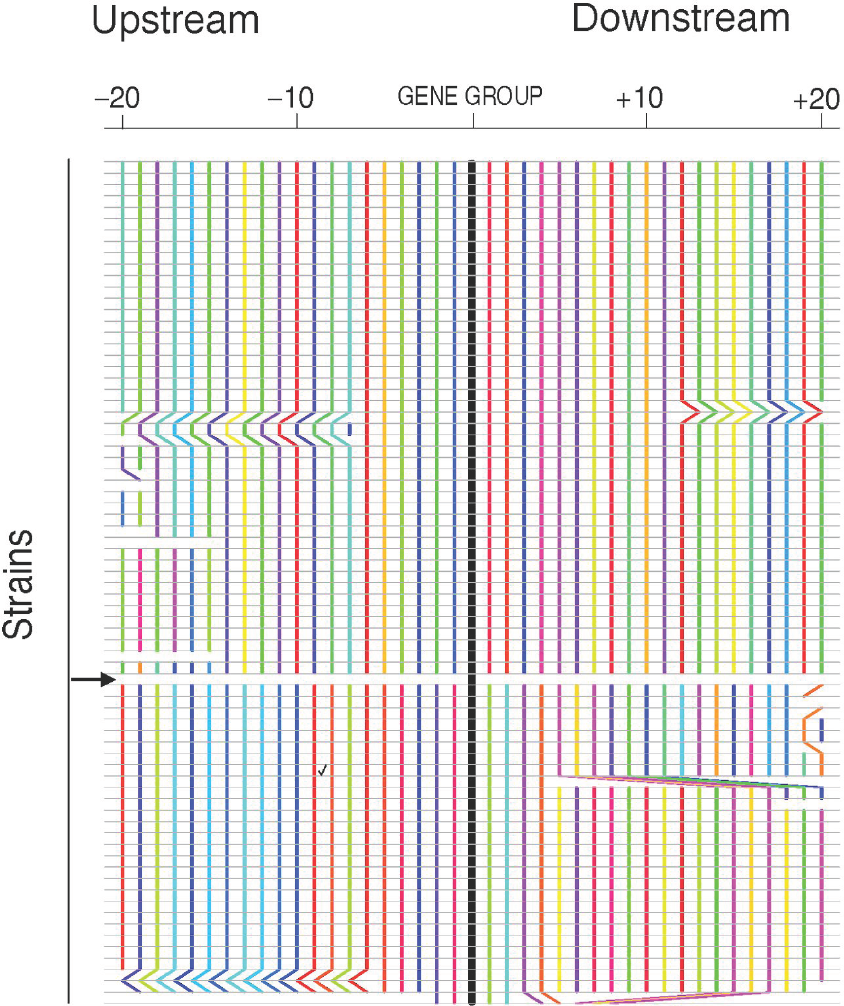
Synteny visualisation. Horizontally there are 41 gene segments depicted from each strain in the gene group. 20 genes upstream, and 20 genes downstream of the gene in the gene group being investigated (thick black vertical line). Each coloured vertical line traces the placement of one gene shared between different strains. A gap signifies that the gene in question is not present in the two strain on either side of the gap. A shift in position of a coloured line denotes a shift in order of the given gene in one strain compared to the other. The middle black vertical line traces the genes from the gene gene group whose synteny is being visualised, while the 20 coloured lines to each side trace genes located immediately up and downstream. Due to the number of unique genes and random assignment of colours some may look similar without actually being the same. The black arrow denotes the point of point of separation between two syntenic clusters within the same gene group.

In order to validate the hypothesis behind the co-ortholog aware disambiguation algorithm two things were needed. First, demonstrating the existence of poor synteny between co-orthologs in a gene group, when compared to orthologs in the same group was required. This would verify that the genomic neighbourhood of duplicated genes differ greatly.

In order to demonstrate this, each group containing co-orthologs was split into two subgroups, one with all the orthologs, and the other containing all co-orthologs.

[A B1 B2 B3 C D E1 E2 F G] *(Synteny 21.1)* =>

**Orthologs:**[A C D F G] *(Synteny 25.4)* **Co-orthologs:**[B1 B2 B3 E1 E2] *(Synteny 16.1)*

Illustrated in the example above is an instance where the subgroup consisting of orthologs has a higher internal synteny than the subgroup with the co-orthologous genes. The low synteny among co-orthologs reveals that their syntenic neighbourhood is more dissimilar than the neighbourhoods of the orthologs. The large difference in average synteny between genes in the co-orthologous subgroups (**Table 2**) was a strong indicator that the disambiguation algorithm would be able to clearly separate them based on synteny.

The second demonstration needed was to show that among co-orthologs in a gene group one gene, the true syntelog, would have much better synteny with the orthologs in the gene group, identifying it as the original copy, while revealing the other co-orthologs to be paralogs. In order to test this hypothesis the synteny score of the co-orthologous gene that was accepted as the true syntelog, and the other co-ortholog(s) that were rejected by the disambiguation algorithm were plotted in separate box plots (**Figure 5**). As expected, the median synteny score of the rejected genes were much lower than that of the genes accepted as true syntelogs.

**Figure 5.**
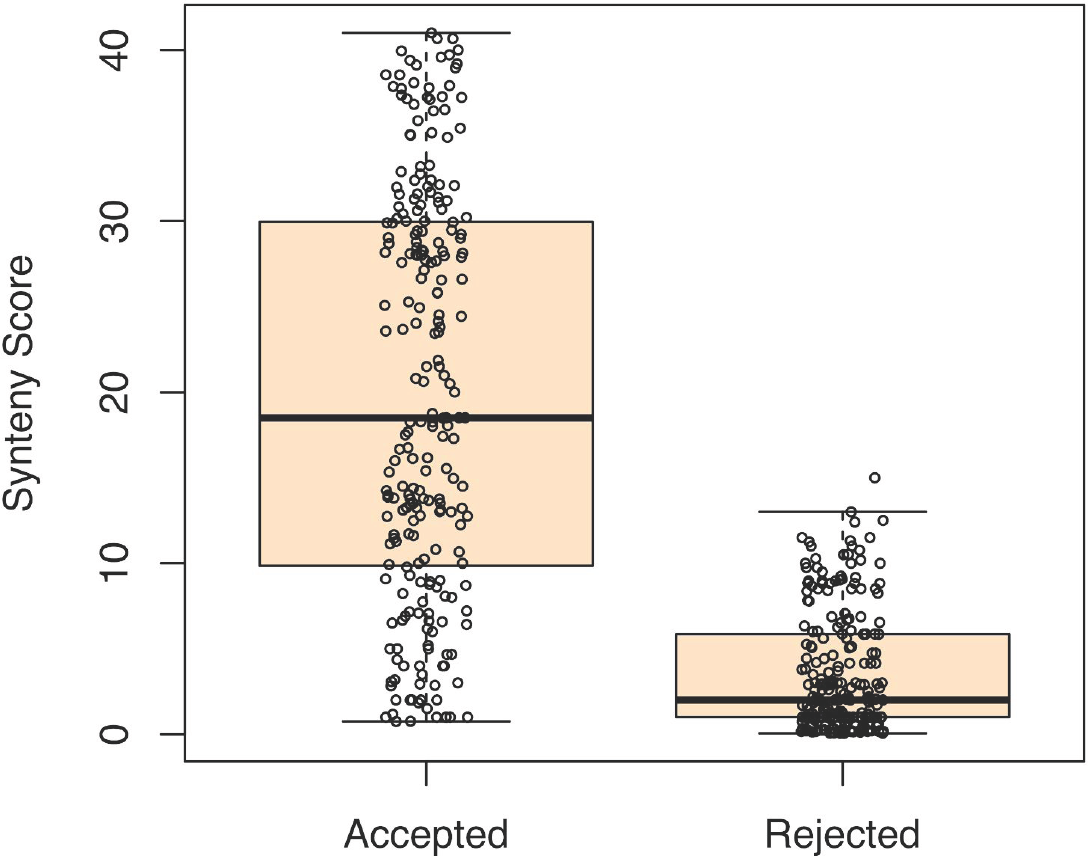
Synteny score comparison of co-orthologs from the same strain being scored against the orthologous genes of the group. “Accepted”: synteny scores of co-orthologs that scored the highest and were accepted as the true syntelog. “Rejected”: scores of the other candidates which scored lower and were rejected. The median and spread of rejected paralogs are much lower than the true syntelogs.

During disambiguation many orphaned genes were being produced by a small subset of strains, when on average the same number of orphans were expected produced by all strains (**Figure 6**).

**Figure 6.**
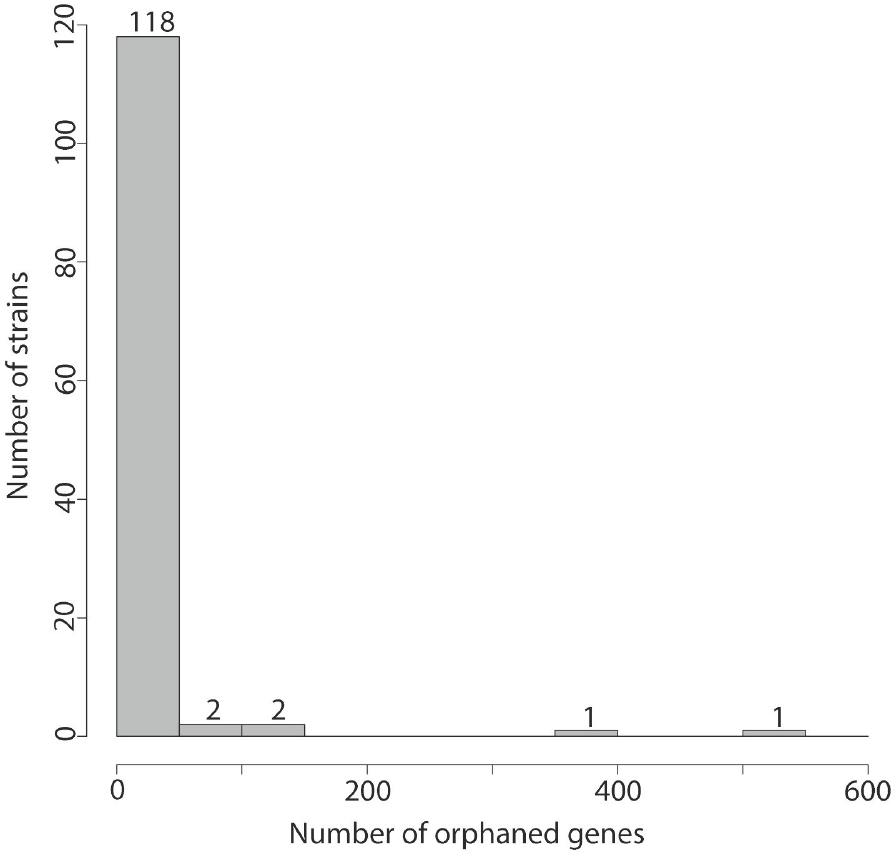
Histogram of the number of strains with a given number of orphaned genes. Most of the strains have under 50 orphaned genes, but a small subset of strains have substantially more.

To investigate the cause, a visualization tool capable of displaying genome-wide synteny scores, GC3s levels, and paralog relationships was built into S3K. This revealed large duplicated regions as the cause of the large number of orphan genes from certain strains (**Figure 7**).

**Figure 7.**
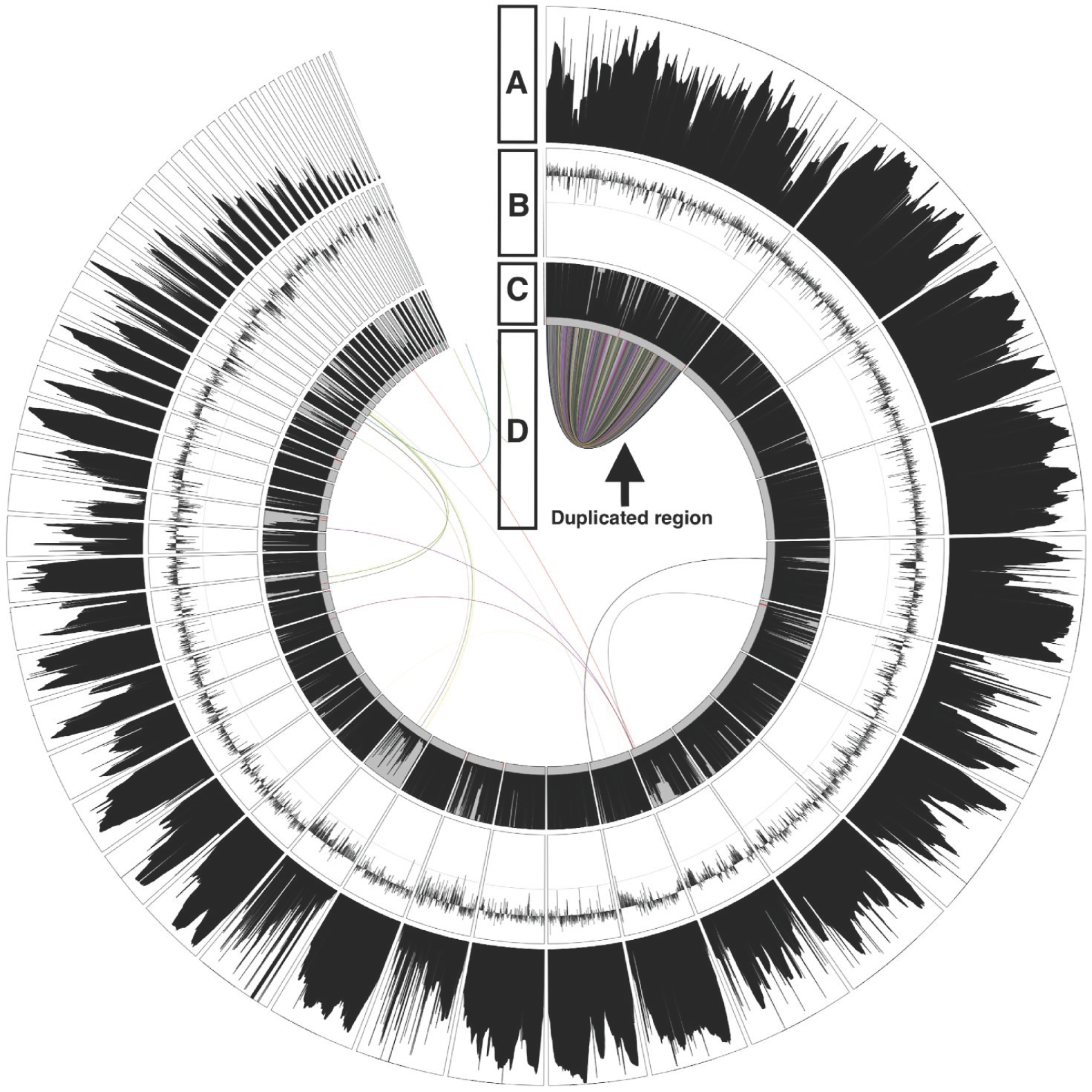
Charting the genome of a strain with an unusually large number of orphaned genes produced during disambiguation revealed a large duplicated region (pointed out by the arrow) as the root cause. **A)** Height map of gene synteny. **B)** GC3s chart with the middle grey line denoting 50%. **C)** Height map of the abundance of each gene across all strains. **D)** Map of co-orthologs where each coloured line connects a co-ortholog pair.

## 5 Discussion

Many techniques to detect paralogs have been attempted, such as using phylogenies [Kim 2008], BLAST scores [Moreno-Hagelsieb 2007], and even synteny [Lechner 2014]. Many of these techniques are mostly concerned with establishing correct phylogenies among distantly related species, not detecting true syntelogs. The software with the closest approach to S3K is the PoFF extension of ProteinOrtho, which enables formation of gene groups with synteny as a weighed factor during cluster disentanglement [Lechner 2011]. However, even gene groups generated using PoFF will contain co-orthologs. Thus, the need for retaining a single copy of a gene from each strain, this being the true syntelog, in each gene group has not been addressed by any of the aforementioned techniques. In order to best serve this need an algorithm was developed based on the ability to precisely measure synteny between two genes.

Synteny analysis of genomes has been employed in studies involving orthology verification [Frantzeskakis 2018], examining chromosomal reconstruction using synteny blocks in distantly related species such as humans and mice [Singa 2007], and detection of horizontal gene transfer [Adato 2015]. Until S3K, no synteny based tools capable of assigning a numerical value to the degree of synteny between two genes, and use this measurement to remove paralogs from gene groups has been made available for general research use.

The large difference in synteny measured between co-orthologous and orthologous pairs (**Table 2**) supports the hypothesis that synteny can be used to distinguish and pick out the true syntelog between co-orthologous pairs. Visualisation of the synteny scores for the co-ortholog, which was accepted as the true syntelog, and the rejected competing candidates (**Figure 5**) reveals a clear difference in median and spread of scores, which further supports that an intact genetic neighbourhood is a valid way of determining true syntelogs, at least in relatively closely related bacterial strains.

The average synteny between orthologs was approximately 20 in our dataset, so the S3K tool was built with a neighbourhood size [n] of 40, twice the average, and five times the average for co-ortholog pairs (Table 2), in order to leave plenty of room for catching potential edge cases.

Larger neighbourhood sizes could have been chosen, [n]=100, 200 or even setting [n] to the size of the contigs. But looking too far to each side would remove focus of the score away from the immediate neighbourhood of the two genes being compared and could result in misleading scores.

One potential problem with our dataset was that with a standard neighbourhood size of 40 there were 3,155 contigs (48%) which contained fewer than 41 genes (Table 1). These could potentially negatively impact synteny score calculations and result in artificially lowered scores since genes residing in them could never achieve a perfect synteny score of 40 as there were not enough genes in their contig neighbourhood.

This could be solved in one of two ways, either lowering the neighbourhood size significantly, and risk potentially lowering the fidelity and range of synteny scores, or improve the assemblies themselves so they were less fragmented.

However, closer inspection revealed that only about 2% of total dataset genes resided on short contigs (Table 1). So a neighbourhood size of 40 was kept to benefit synteny analysis of about 98% of the genes that were part of larger contigs.

In addition [n]=40 should be large enough to meet the needs of other datasets as well, since 40 genes is likely to cover a large enough area to allow accurate true syntelog detection in any genome.

One interesting metric produced by ProteinOrtho is the connectivity measure for each of the generated gene groups. During its disentanglement process connectivity, which is a number in the range 0 to 1, is what ProteinOrtho uses to measure how well the genes in a cluster fit in with one another [Lecher 2011]. A connectivity of 0.1 or lower is the threshold ProteinOrtho uses to split up a cluster. Connectivity is calculated based on sequence similarity, unless the PoFF extension is enabled, in which case synteny will account for half of the connectivity score. So in a sense the connectivity score produced by ProteinOrtho, with PoFF disabled, is not unlike the synteny score that we assign to each group, only it is based on gene sequence similarity, rather than synteny.

Due to the similar nature of both measures it was of interest to see whether there was a good correlation between gene the group connectivity measure, and the average synteny score for each group.

However charting group synteny against group connectivity (**Figure 8**) did not reveal a strict linear relationship.

**Figure 8.**
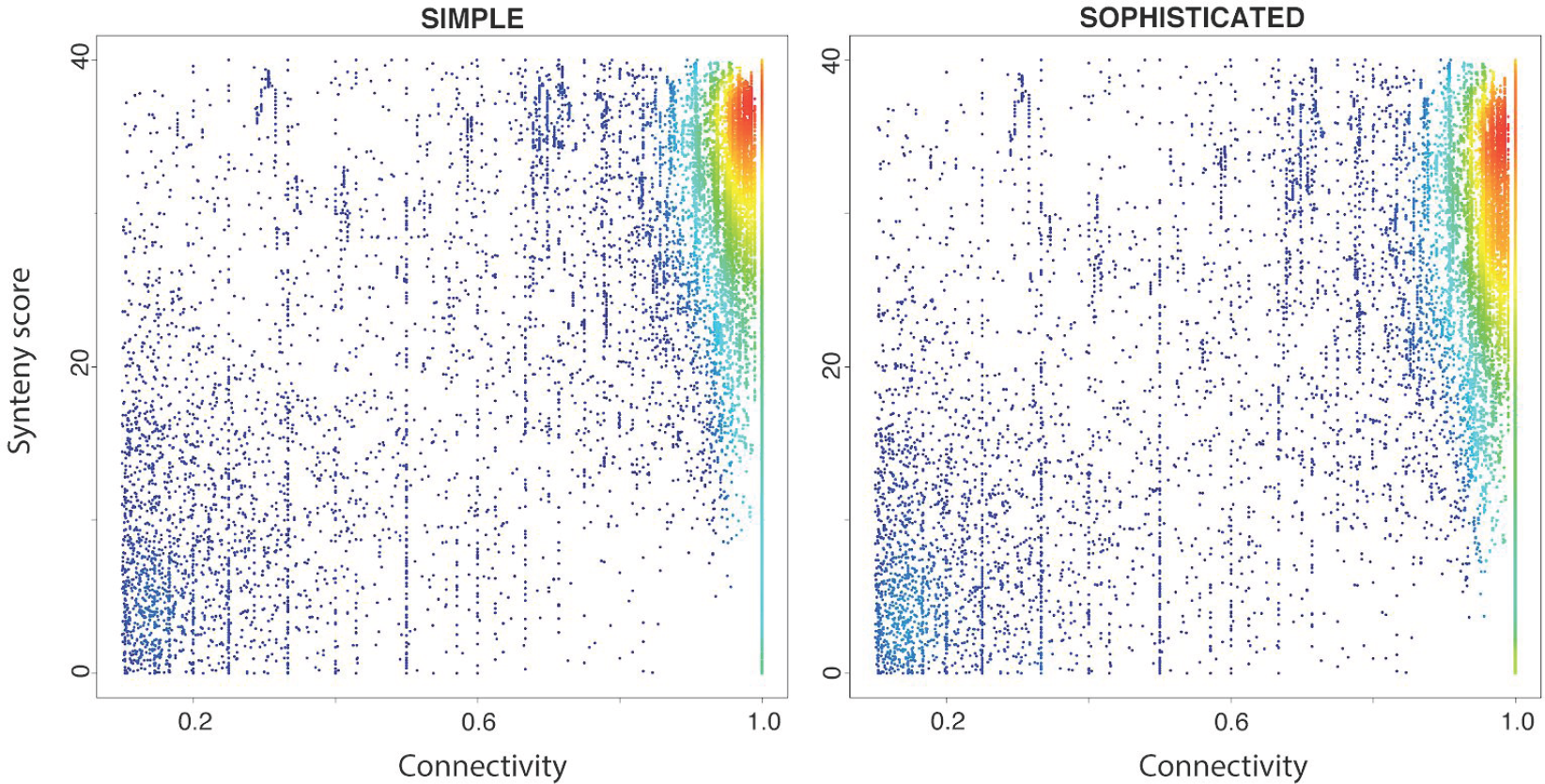
Group SOPHISTICATED OR SIMPLE synteny scores charted on y-axis against the group connectivity measure on x-axis does not reveal a strict linear relationship.

A strong clustering in the top right corner for both SOPHISTICATED AND SIMPLE synteny scores do show good correlation for groups with high scores in both category. A clustering in the lower left corner reveals that low synteny and connectivity scoring groups often overlap.

One interpretation of this is that groups with very similar nucleotide sequences also have very similar neighbourhoods. The smaller cluster near the bottom left suggests that the groups placed there contain many genes that perhaps are not truly homologous, and do not belong together.

Such groups are possible candidates for being split up by removing genes with low sequence and syntenic similarity with other genes in the group into their own separate group.

In such scenarios the *“--splitgroups”* option in S3K can be toggled to utilize an algorithm that splits gene groups based on the presence of syntenic clusters within them (Figure 4). However, in practice such syntenic clusters were proven to be quite prevalent in our dataset, making automated group splitting inadvisable as it would lead to high fragmentation of larger gene groups. As such this experimental functionality is best used manually on groups where splitting based on synteny clusters is explicitly desired.

## 6 Conclusion

During the course of development many novel uses for syntenic analysis, and genome visualisation were developed and proved useful in large-scale analysis of bacterial genomes. At a very basic usage level S3K was an easy to use and versatile tool for assessing and quickly gaining a overview of bacterial populations and act as a robust foundation and first step towards further downstream analysis.

The large discrepancy between synteny scores of the paralog selected as the true syntelog, and the other candidates is a strong indicator of the viability of our synteny based disambiguation method for our dataset. Although this does not guarantee that the method is viable for any dataset, the statistics and charts produced by the S3K will aid in making such an assessment on a dataset by dataset basis.

We are making S3K available for use in hope that it might ease synteny analysis and bacterial genome visualisation.

## 7 Acknowledgements

This work was funded by grant no. 4105-00007A from Innovation Fund Denmark (S.U.A.).

## 8 Author contributions

Conceptualization: CM, SUA; Methodology: CM; Software: CM; Validation: CKS, SM, CM; Formal analysis: CM; Investigation: CM; Resources: SUA; Data curation: CM; Writing - original draft: CM; Writing - review and editing: CM, CKS, SM, SUA; Visualisation: CKS, CM, SM; Supervision: SUA; Project administration: SUA; Funding acquisition: SUA.

